# Compounds derived from *N,N*-dimethyldithiocarbamate are effective copper-dependent antimicrobials against *Streptococcus pneumoniae*

**DOI:** 10.1101/2022.09.23.509297

**Authors:** Sanjay V. Menghani, Yamil Sanchez-Rosario, Chansorena Pok, Renshuai Liu, Feng Gao, Henrik O’Brien, Miranda J. Neubert, Klariza Ochoa, Riley D. Hellinger, Wei Wang, Michael D. L. Johnson

**Author notes:** Corresponding Author: Michael D. L. Johnson, University of Arizona, 1656 E. Mabel St. / P.O. Box 245221 / MRB 213 (office), Tucson, AZ 85724.

## Abstract

*N,N*-dimethyldithiocarbamate (DMDC) is a potent copper-dependent antimicrobial against several pathogens, including *Streptococcus pneumoniae*. Despite the availability of several vaccines against multiple disease-causing strains of *S. pneumoniae*, the rise of antimicrobial resistance and pneumococcal disease caused by strains not covered by the vaccine creates a need for developing novel antimicrobial strategies. We derived novel compounds from DMDC and tested their effectiveness as copper-dependent antimicrobials against *S. pneumoniae* through *in vitro* growth and killing curves. Compounds that caused a growth defect and were bactericidal *in vitro* were tested against other strains of *S. pneumoniae* and in complex with different transition metals. We found two compounds, sodium *N*-benzyl-*N*-methyldithiocarbamate and sodium *N*-allyl-*N*-methyldithiocarbamate (herein “Compound 3” and “Compound 4”), were effective against TIGR4, D39, and ATCC® 6303™ (a type 3 capsular strain) and further increased the internal concentrations of copper to the same previously reported levels as with DMDC and copper treatment. We found that both Compound 3 and Compound 4 were bacteriostatic in combination with zinc. We tested Compound 3 and Compound 4 *in vivo* against a murine pneumonia model, finding that Compound 3, and not Compound 4, was effective in significantly decreasing the bacterial burden in the blood and lungs of *S. pneumoniae*-infected mice. We found that the combination of Compound 3 and copper made the pneumococcus more susceptible to activated macrophage mediated killing via an *in vitro* macrophage killing assay. Collectively, we demonstrate that derivatizing DMDC holds promise as potent bactericidal antibiotics against *S. pneumoniae*.

## Introduction

*Streptococcus pneumoniae* (also known as the pneumococcus) is a Gram-positive bacterium that is a leading cause of pneumonia, otitis media, meningitis, and sepsis. As such, the pneumococcus causes a significant disease burden in the pediatric and elderly populations despite effective vaccines against the most common strains (1). Normally a commensal of the nasopharynx and upper respiratory tract, the causes and gene regulation of pathogenic transformation is an ongoing topic of investigation (2). In addition, the United States Centers for Disease Control and Prevention (CDC) has identified drug-resistant *S. pneumoniae* as a serious threat in their 2019 report “Antibiotic Resistance Threats in the United States,” with approximately 30% of clinical strains of *S. pneumoniae* displaying resistance to at least one antibiotic and many more harboring multidrug resistance (3).

The global rise of antibiotic-resistant bacterial strains has created a renewed desire to develop novel antimicrobials. Traditionally, most antibiotics used clinically against *Streptococcus pneumoniae* have targeted a single bacterial protein or enzyme for bactericidal or bacteriostatic effect, like the bacterial cell wall for β-lactams or bacterial RNA translation for aminoglycosides (4, 5). Unsurprisingly, the global selective pressure of using single-target antibiotics allows bacteria to escape antibiotic coverage by acquiring fewer mutations. Clinically relevant resistant strains are seen within years of the introduction of novel antibiotics, as was observed for penicillin and methicillin resistance in the bacteria *Staphylococcus aureus* in the mid-1940s (6, 7). Beyond the less than 25 bacterial proteins and enzymes inhibited by these traditional antibiotics, computational studies have identified 300 essential and highly conserved antimicrobial targets to advance drug discovery (8, 9). A drug targeting multiple bacterial enzymes at once would require a bacterium to acquire multiple mutations to develop antibiotic resistance (10). Another strategy for developing novel antimicrobials is drug repurposing, testing the *in vitro* antimicrobial activity of molecules or drugs that have United States Food and Drug Administration (FDA) approval for other indications (11). The recent identification of antimicrobial properties of the drug disulfiram has combined these two principles (12-15).

The drug disulfiram (also known as tetraethylthiuram disulfide, TETD, or the brand name Antabuse) has been used to treat alcohol dependence for decades, despite a significant decrease in clinical use over time. In the body, disulfiram is rapidly converted to its metabolite diethyldithiocarbamate (DETDC), which is complexed with divalent metal zinc and copper ions (16). Recently, there has been a growing body of evidence for the antimicrobial properties of disulfiram’s metabolites. In 2020, a group led by Ghosh *et al*. identified Zn^2+^-diethyldithiocarbamate as a potent antiparasitic agent against *Entamoeba histolytica* and other parasites (17). In 2021, our group performed a targeted antimicrobial drug screen involving dithiocarbamate compounds (18). In that study, diethyldithiocarbamate (DEDC) and *N,N*-dimethyldithiocarbamate (DMDC) were investigated, with DMDC being identified as a potent copper-dependent antimicrobial against *S. pneumoniae, Coccidioides immitis*, and *Schistosoma mansoni* (18). Since this study, other groups have also identified copper-dependent antibiotic potential for dithiocarbamates (19).

Mechanistically, DMDC significantly increases the intrabacterial concentration of copper, putting stress on the bacteria to export copper before mismetallation inactivates or reduces the efficiency of key bacterial enzymes, eventually leading to bacterial death (20, 21). Further, when the excess oxidized copper enters the bacteria, it diminishes the reductive capacity of the bacteria, which must reduce copper ions in order to export them (21-26). As a result, the pneumococcus cannibalized its capsule, making macrophage mediating killing more efficient, and presumably to extract electrons from the sugar building blocks to maintain the intrabacterial reducing environment (21, 27).

In this study, we derived dithiocarbamate compounds from the starting compound, DMDC. We tested the *in vitro* copper-dependent antimicrobial efficacy of these compounds (named “Compound 1” through “Compound 5”) against *S. pneumoniae* TIGR4, D39, and Type 3 strains. The zinc-dependent toxicity of these compounds was also tested against TIGR4 and found to be bacteriostatic. Next, we tested the *in vivo* efficacy of these compounds in a murine pneumonia model determining they reduced the bacterial burden in the lungs. Finally, we tested the ability of these compounds to enhance *in vitro* macrophage-mediated killing, finding the combination of Compound 3 and copper to be cleared more efficiently. Overall, this study identifies novel compounds in the dithiocarbamate class that are broadly anti-pneumococcal *in vitro* and effective *in vivo*.

## Results

We derived DMDC into five compounds that retain putative copper ionophore properties (Table 1). The six derived compounds were named “Compound 1” through “Compound 5”. Each compound was initially screened by performing *in vitro* growth curves to determine if a growth defect was associated with either the compound alone or in combination with copper. Next, successful compounds were tested in killing curves to determine if there was a bactericidal or bacteriostatic effect of copper, compound, or combination treatment. Compound 3 and Compound 4 showed statistically significant growth defects for the combination treatments in comparison to an untreated control in TIGR4 *S. pneumoniae* (Figure 1A and 1C). Compound 1, Compound 2, and Compound 5 had no effect, and thus, experiments with these compounds were discontinued (data not shown). Further, both Compound 3 and Compound 4 showed significant bactericidal activity in killing curves (Figure 1B and 1D). From these data, we concluded that Compound 3 and Compound 4 are viable copper-dependent bactericidal antibiotics that warrant further investigation.

**Table 1.**
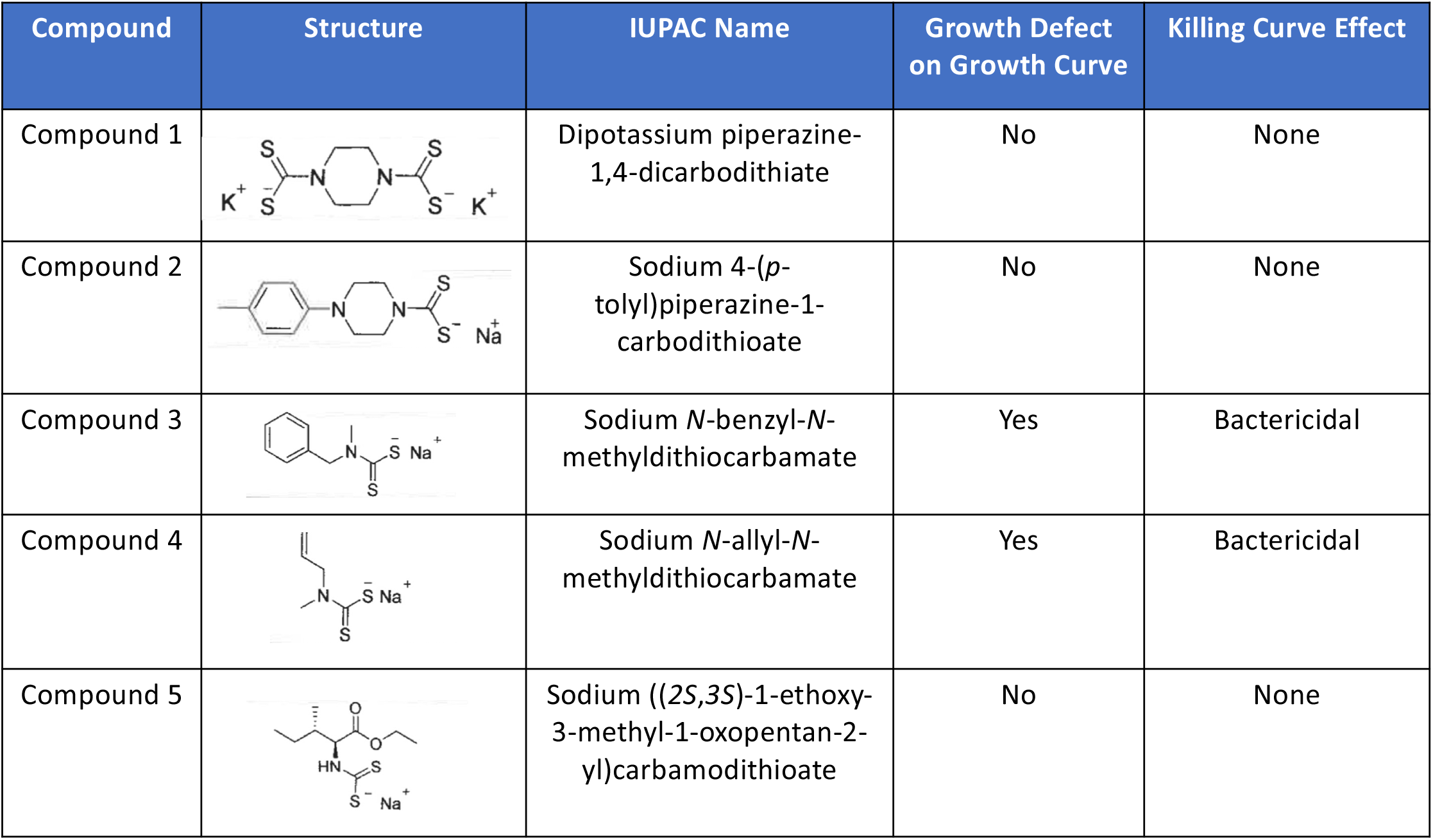
Summary of screening results of each compound derived from DMDC. Structure of each compound, growth curve assay results, and killing curve assay results are summarized for each compound.

| Compound | Structure | IUPAC Name | Growth Defect on Growth Curve | Killing Curve Effect |
| --- | --- | --- | --- | --- |
| Compound 1 |  | Dipotassium piperazine-1,4-dicarbodithiate | No | None |
| Compound 2 |  | Sodium 4-(*p*-tolyl)piperazine-1-carbodithioate | No | None |
| Compound 3 |  | Sodium *N*-benzyl-*N*-methyldithiocarbamate | Yes | Bactericidal |
| Compound 4 |  | Sodium *N*-allyl-*N*-methyldithiocarbamate | Yes | Bactericidal |
| Compound 5 |  | Sodium ((*2S,3S*)-1-ethoxy-3-methyl-1-oxopentan-2-yl)carbamodithioate | No | None |

**Figure 1.**
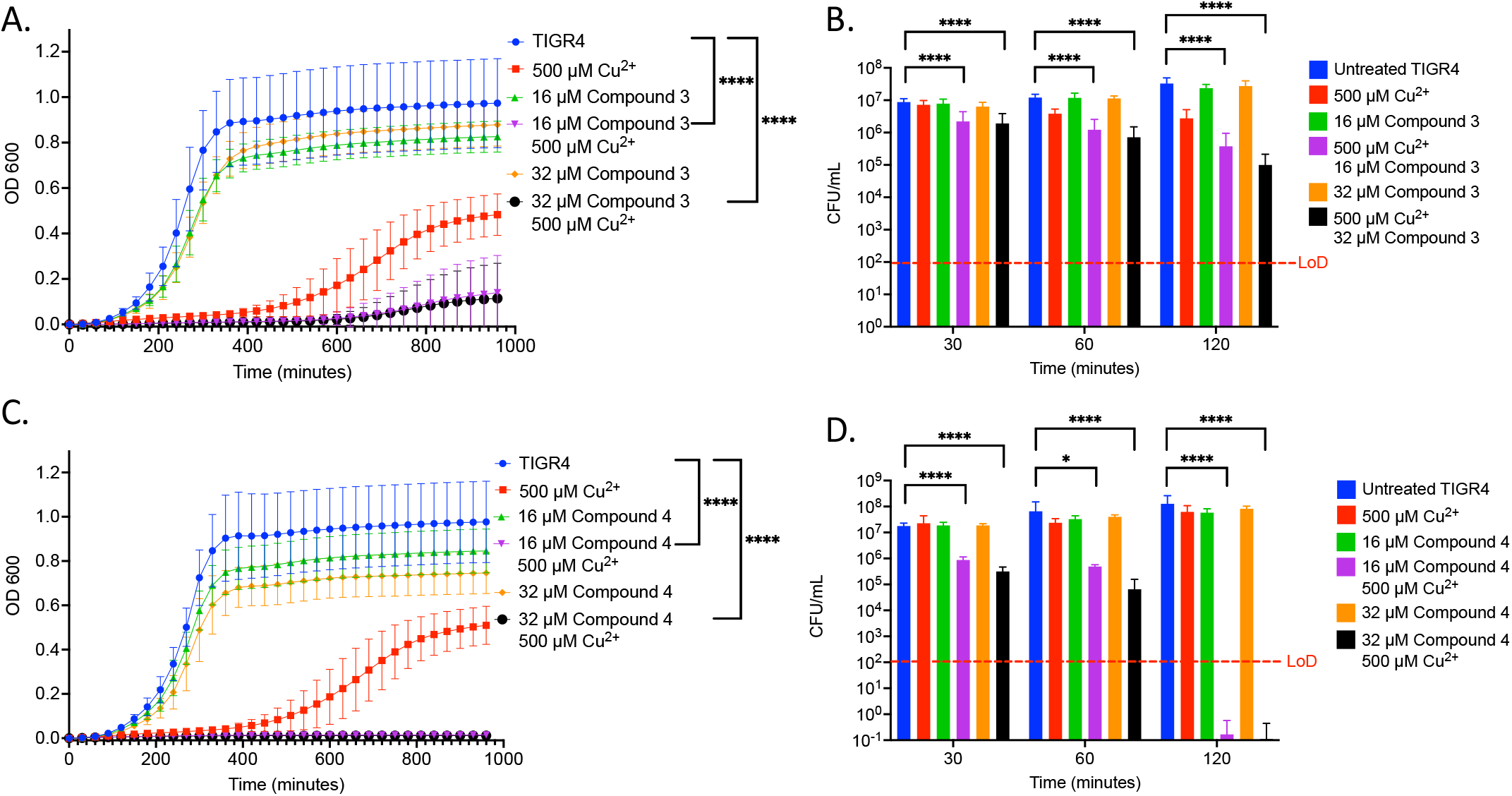
*In vitro* testing of copper-dependent antimicrobial activity of dithiocarbamate compounds derived from *N,N*-dimethyldithiocarbamate (DMDC) (A) Growth curve assay of WT TIGR4 exposed to the indicated concentrations of copper sulfate and Compound 3. (B) Killing curve assay of WT TIGR4 starting with an inoculum of 5.6×10^6^ CFU/mL in M17 media supplemented with indicated concentrations of copper sulfate and/or Compound 3. (C) Growth curve assay of WT TIGR4 exposed to the indicated concentrations of copper sulfate and Compound 4. (D) Killing curve assay of WT TIGR4 starting with an inoculum of 1.3×10^6^ CFU/mL in M17 media supplemented with indicated concentrations of copper sulfate and/or Compound 4. All bars indicate mean ± SD with n = 12-18 across 3 independent replicates for growth curves and n = 9 across 3 independent replicates for killing curves. Statistical differences were measured by one way ANOVA; ns = not significant, *p < 0.05, **p < 0.01, ***p < 0.001, and ****p < 0.0001.

We next examined the ability of the compounds to increase the concentration of copper inside TIGR4. Using DMDC plus copper, we previously observed a significant increase in internal copper relative to copper or DMDC treatments alone (21). We used Inductively Coupled Plasma Optical Emission Spectroscopy (ICP-OES) for this measurement. Compound 3 and Compound 4 showed significant increases in the combination treatment relative to the compound or copper alone (Figure 2A-D). Thus, Compound 3 and Compound 4 facilitate copper entry into the bacteria.

**Figure 2.**
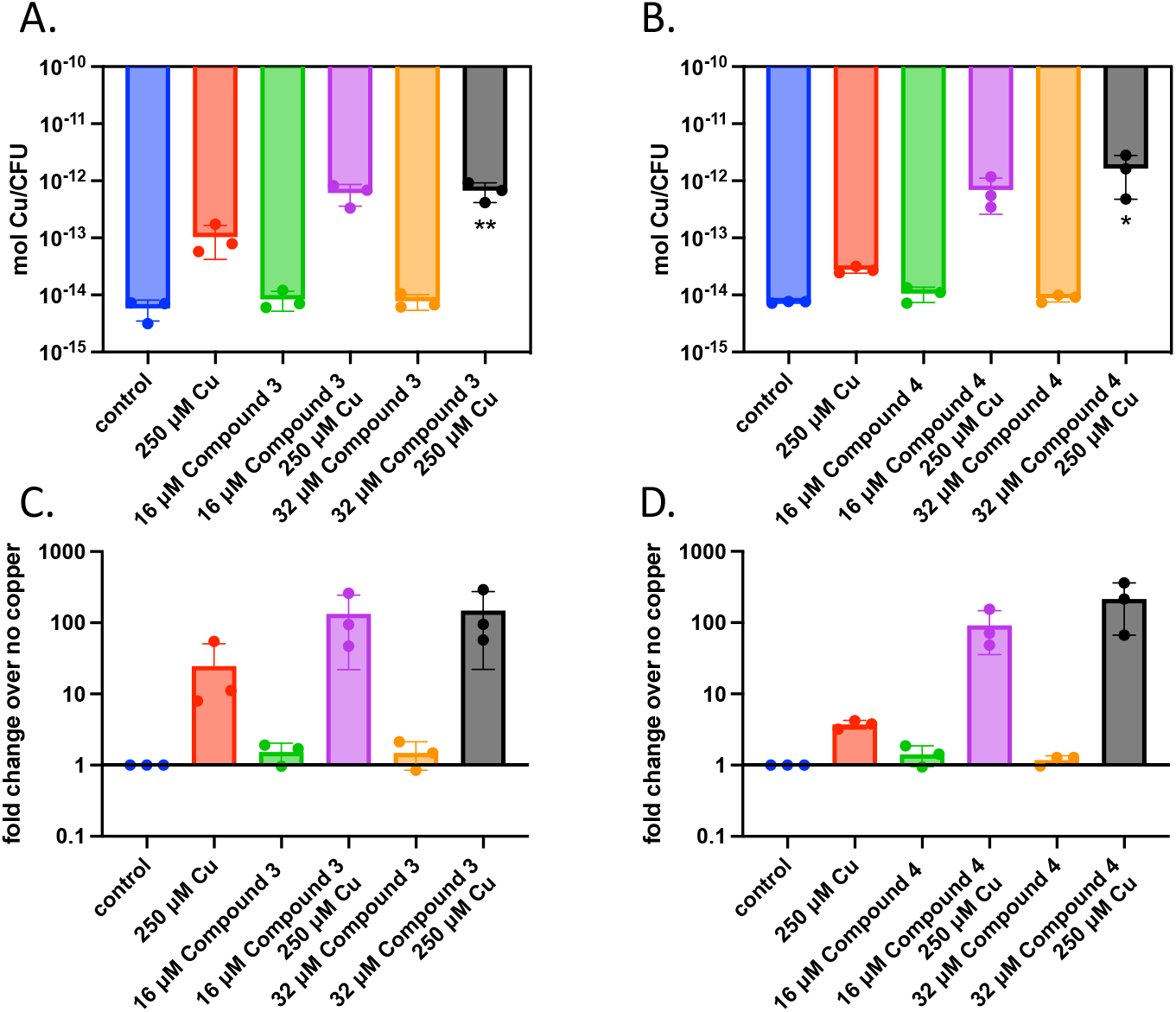
Intracellular copper concentration of TIGR4 after treatment with compounds and copper. Exponentially growing bacteria were exposed to a combination of compound and copper for 30 minutes. ICP-OES was used to measure intracellular copper. (A) and (B) represent moles of copper per CFU for Compounds 3 and 4 while (C) and (D) represent the fold change over the parent strain with nothing added. Experiments were performed in triplicate with statistical significance determined by an Ordinary one-way ANOVA; *p < 0.05, **p < 0.01.

Determining copper affinity is notoriously difficult due to relatively high-affinity values. Thus, we sought to at least compare the binding rates of the compounds to DMDC using DsRed2, a protein that has its fluorescence quenched by copper. The stoichiometry of DMDC to copper is 2:1 (28); this premise was used to provide the compounds in excess for the chelation assays. Using our compounds to compete for copper to reverse the quenching of copper to DsRed2, we found initial hierarchies in their ability to chelate copper and restore fluorescence (Figure 3). Interestingly, bactericidal compounds Compound 3 and Compound 4 exhibited more prolonged periods to quench DSRED2 as compared to parent compound DMDC. Thus, while relative affinities and compound-mediated copper influx could play a role in the overall effectiveness of the antibiotic capacity of the compound, this process is not linear and likely highly dependent on the microbe-of-study’s phenotype and ability to flux metal in and out of the organism.

**Figure 3.**
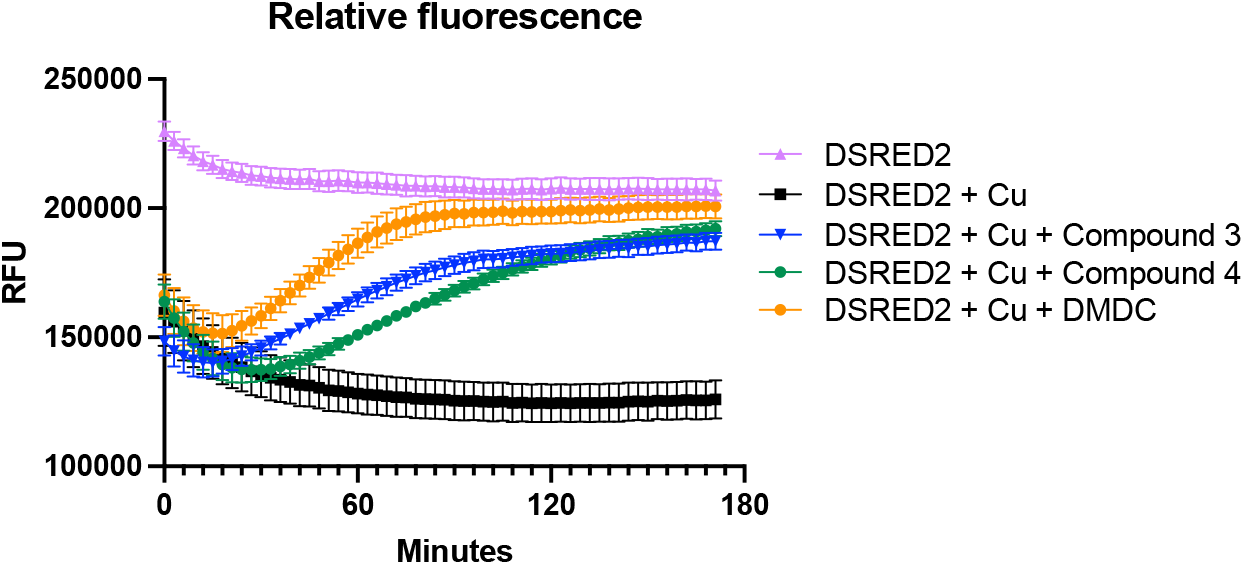
Chelation ability of compounds and DMDC for copper relative to DsRed2. Restoration of DsRed2 fluorescence treated with 100 µM copper and the different chelating compounds was measured and compared to a DsRed2-only control. Compounds were added at 400 µM. The graph shows the standard error of the mean of three independent experiments.

In our group’s targeted small-molecule screen that identified DMDC as a potent copper-dependent antibiotic, we investigated combinations of many divalent metals with DMDC to determine if other metals can display toxicity (18). To test the zinc-dependent toxicity, we performed killing curves using TIGR4 against combinations of zinc, Compound 3 or Compound 4, and combinations of compound + zinc. We observed that the combination of 500 µM Zn^2+^ + 16 µM Compound 3 is bacteriostatic (Figure 4A). Further, we observed that adding 500 µM Mn^2+^ does not rescue the copper-dependent toxicity seen with 250 µM Cu^2+^ + 32 µM Compound 3 (Figure 4B). We repeated these experiments for Compound 4 (Figures 4C and D). While we found that 500 µM Zn^2+^ + 16 µM Compound 4 is bacteriostatic, strikingly, we found that the addition of 500 µM Mn^2+^ does rescue the copper-dependent toxicity seen with 250 µM Cu^2+^ + 32 µM Compound 4 (Figure 4C and D). From these data, we can conclude that Compound 3 and Compound 4 are dithiocarbamate compounds whose copper-dependent toxicity may be mediated by facilitating mismetallation events within the bacterium.

**Figure 4.**
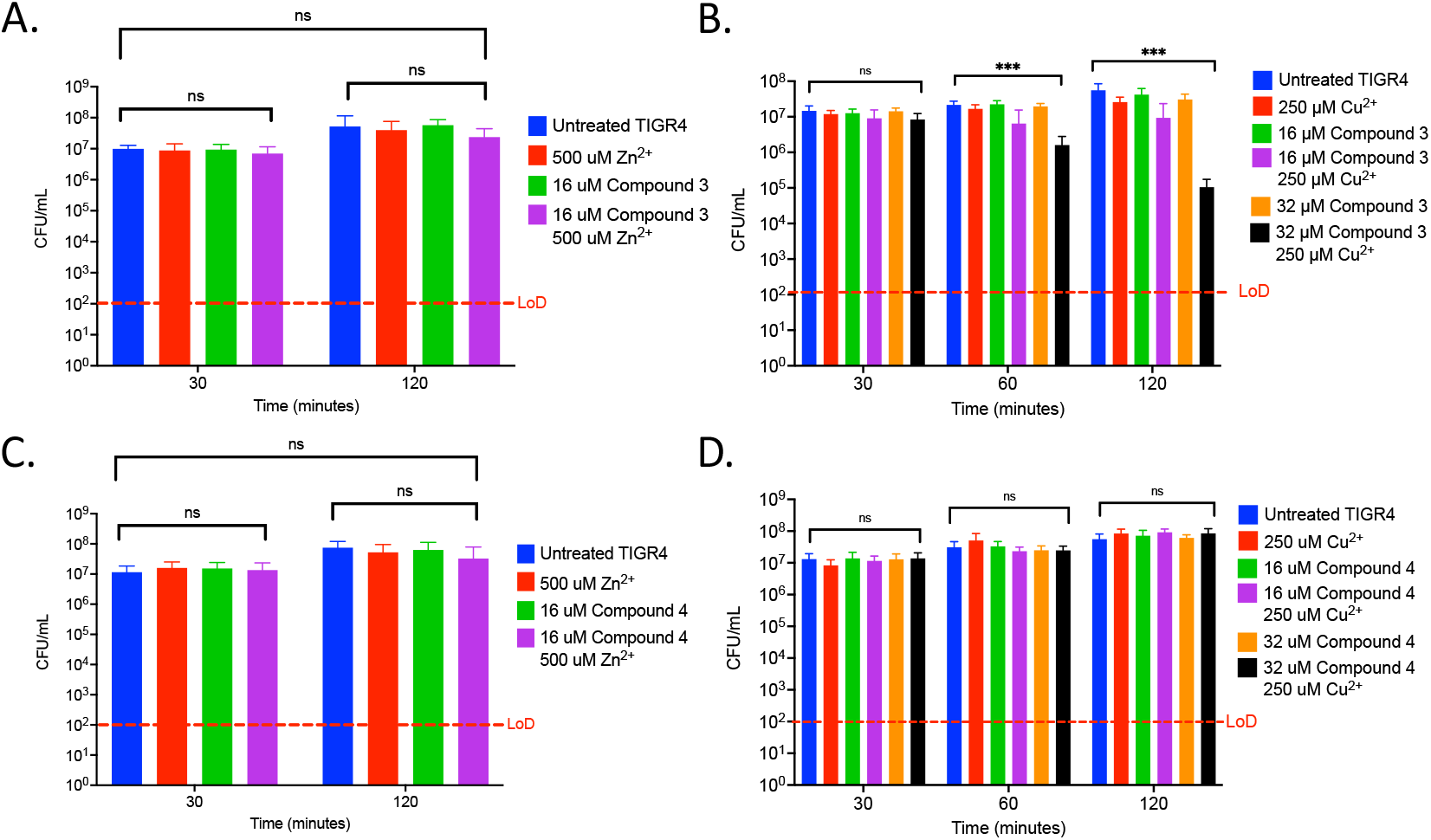
Compound 3 and Compound 4 display zinc-dependent toxicity, and the copper-dependent toxicity of Compound 3 can be rescued by Mn^2+^ supplementation. (A) Killing curve assay of WT TIGR4 strain starting with an inoculum of 9.9×10^6^ CFU/mL in M17 media supplemented with indicated concentrations of zinc sulfate and/or Compound 3. (B) Killing curve assay of WT TIGR4 strain starting with an inoculum of 9.0×10^6^ CFU/mL in M17 media supplemented with 500 µM Mn^2+^ and the indicated copper sulfate concentrations and/or Compound 3. (C) Killing curve assay of WT TIGR4 strain starting with an inoculum of 1.2×10^7^ CFU/mL in M17 media supplemented with indicated concentrations of zinc sulfate and/or Compound 4. (D) Killing curve assay of WT TIGR4 strain starting with an inoculum of 1.1×10^7^ CFU/mL in M17 media supplemented with 500 µM Mn^2+^ and the indicated concentrations of copper sulfate and/or Compound 4. Recovery of bacterial CFU/mL to that of untreated bacteria is seen at t = 120 minutes, with no statistically significant difference between untreated and combination treatment. All bars indicate mean ± SD with n = 9 across 3 independent replicates. Statistical differences were measured by one way ANOVA; ns = not significant, *p < 0.05, **p < 0.01, ***p < 0.001, and ****p < 0.0001.

To directly test if the two compounds are promising antibiotics to progress along the drug discovery pipeline, we tested these compounds in a murine pneumonia model previously used by our lab (18). *S. pneumoniae* TIGR4 bacterial burden is reduced two days post-infection in mice given an intranasal 100% lethal dose (LD_100_). We observed a significant decrease in bacterial titers in the blood and lungs 48 hours post-infection in 8-week-old BALB/c mice given 25 µL of 10 mM Compound 3 (approximately 1.6 mg/kg), but no significant decrease was noted for mice given the same dosage of Compound 4 (Figure 5A-D). Further, as previously observed with DMDC and copper (21), the combination treatment of Compound 3 plus copper also made the pneumococcus more susceptible to activated macrophage mediated killing via a macrophage killing assay as compared to no treatment, copper, or Compound 3 alone (Figure 6). These data show that Compound 3 is a promising antibiotic against *S. pneumoniae* TIGR4 with efficacy *in vivo*.

**Figure 5.**
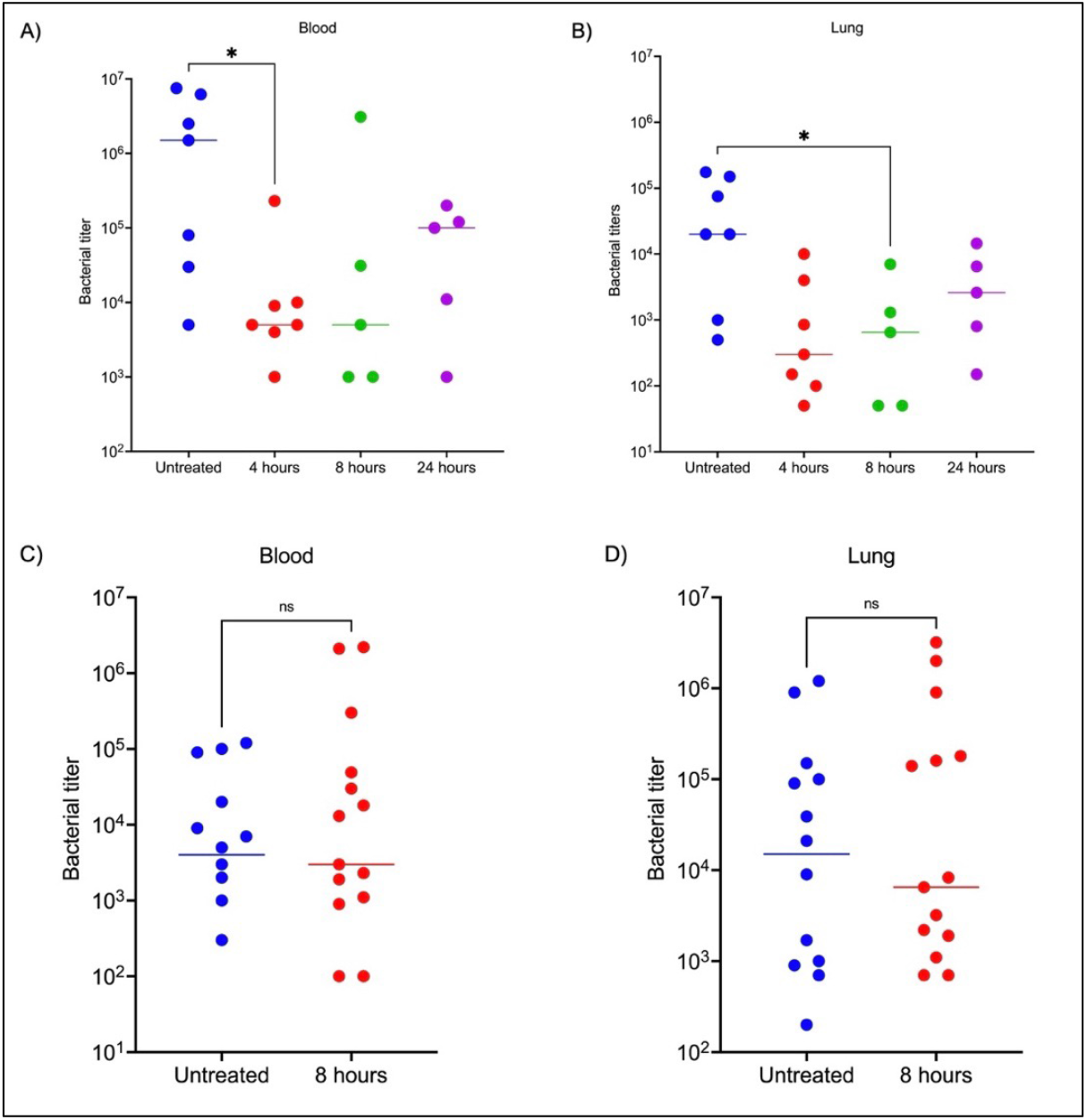
Compound 3 is an effective antibiotic *in vivo* against a murine *Streptococcus pneumoniae* infection model, while Compound 4 is not an effective antibiotic *in vivo*. Groups of 8-week-old female BALB/c mice (n = 5-7) were infected with bacteria at t = 0 and treated with Compound 3 or Compound 4 at 4-, 8-, or 24-hours post-infection or were untreated. At 48 hours post-infection, animals were sacrificed, and blood (A, C) or lung (B, D) bacterial titers were measured. (A) and (B) show Compound 3 treatment reduces bacterial titers in the blood and lungs, respectively. (C) and (D) show that Compound 4 treatment does not reduce bacterial titers in the blood and lungs. Mann-Whitney Wilcoxon rank sum tests were used to measure statistical significance at a P value of <0.05 (*) with no statistical difference noted (ns). The bar within the data set represents the median.

**Figure 6.**
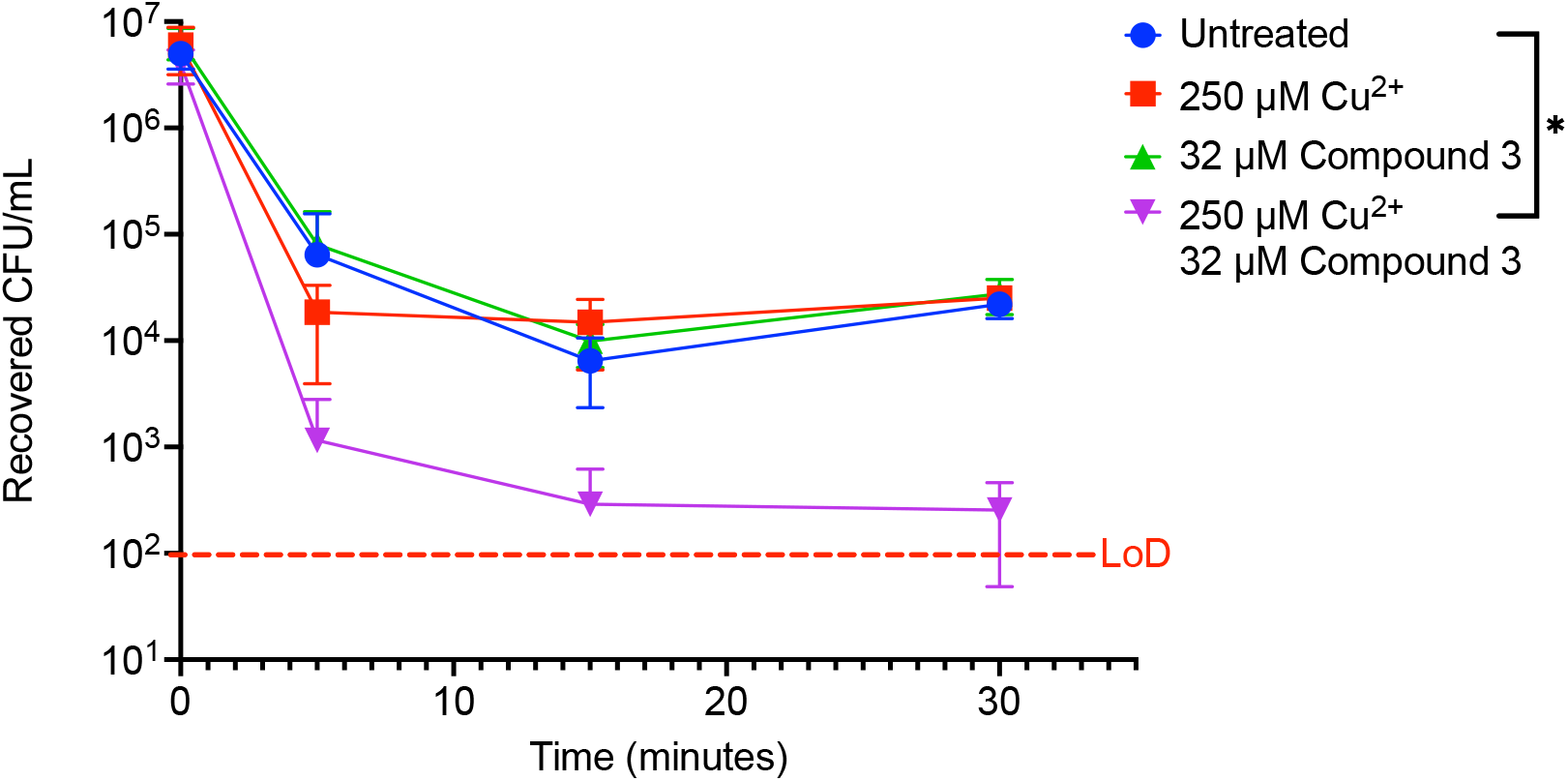
Macrophages display enhanced *post hoc* killing of Compound 3 + Cu^2+^-treated TIGR4 bacteria. WT TIGR4 bacteria that were treated with indicated combinations of Cu^2+^ and DMDC were co-cultured with activated J774A.1 macrophages. The initial inoculum of bacteria added to the macrophages was 7.3×10^6^ CFU/mL (following a 15-minute incubation with indicted conditions), for an MOI of 10. There is a statically significant decrease in recovered CFU/mL between the untreated bacteria and Cu^2+^ + Compound 3-treated bacteria at t = 5 minutes. At this timepoint, combination-treated bacteria were significantly cleared by the macrophages, indicating a rapid *post hoc* bactericidal killing capacity. All bars represent mean ± standard deviation (SD) with n = 12 across 3 independent replicates.

To determine if Compound 3 and Compound 4 are antibiotics that can be widely used against *S. pneumoniae*, we performed killing curves with these compounds against the D39 strain and a type 3 strain (ATCC® 6303™). Compound 3 and Compound 4 were bactericidal in combination with copper against the D39 and the Type 3 strain, and more so (by several logs) than the effects of DMDC versus these strains as seen previously (Figure 7A-D) (18). Lastly, we wanted to measure the intrabacterial concentration of copper with Compounds 3 and 4 against D39 and the type 3 strain. Because we saw increased killing as compared to TIGR4 at the same concentrations (32 µM compound and 500 µM copper), we went lower on our respective concentrations (8 µM Compound and 125 µM copper) to make sure the bacteria were still viable at the time of the measurement. Compound 4 increased copper uptake in D39 and the type 3 strain to a similar degree as with TIGR4 (Figure 8A-D). At these concentrations, only Compound 3 showed increased concentration of copper inside of the type 3 strain, and not D39 (Figure 8A-D). However, not seeing the increase likely has more to do with the concentration used so we could make sure we were comparing viable cells than the lack of effect of the compound. Together with the data from Figure 1, we concluded that Compound 3 and Compound 4 display broad copper-dependent antibacterial effects against several strains of *S. pneumoniae*.

**Figure 7.**
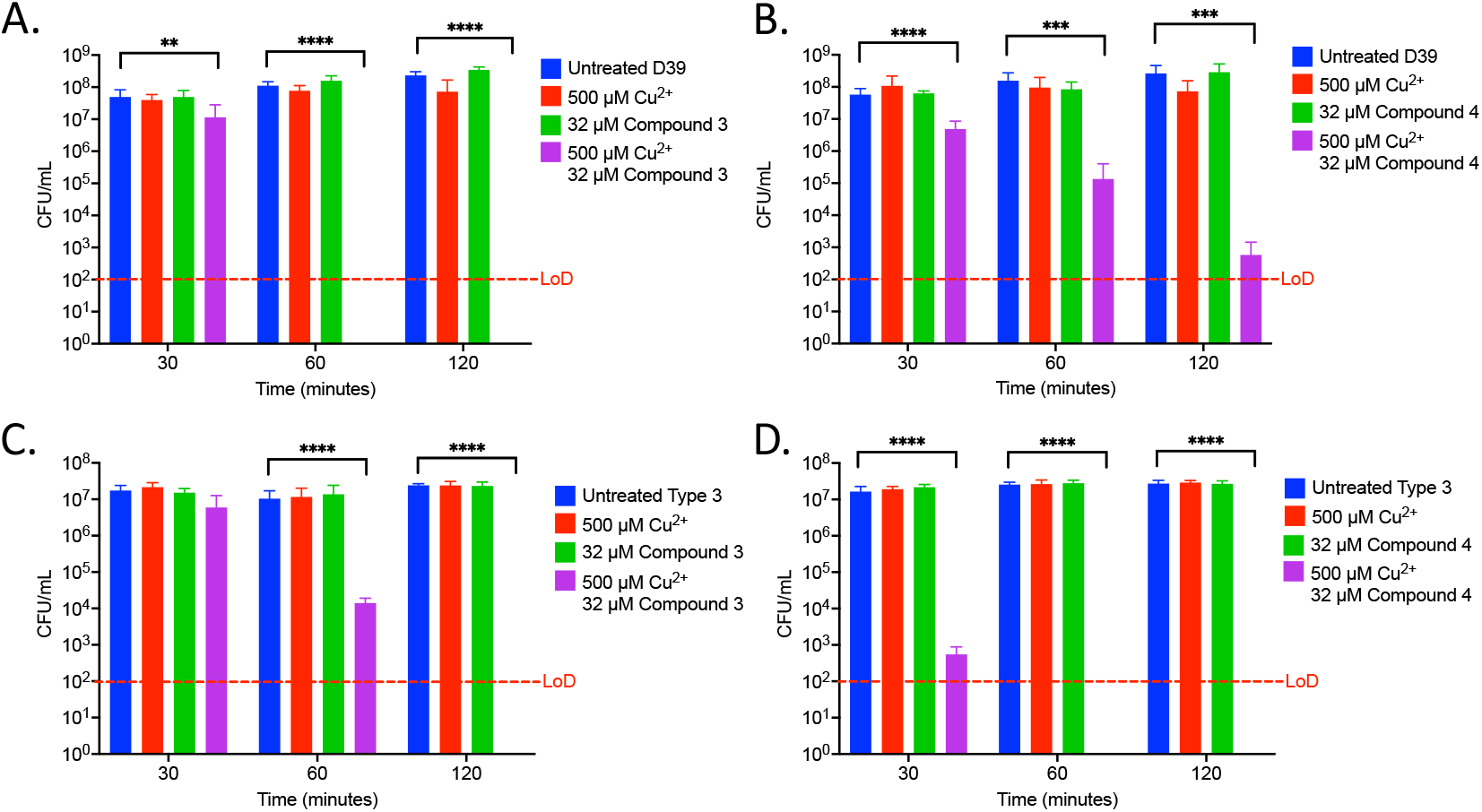
Compound 3 and Compound 4 are broadly antibiotic against multiple species of *S. pneumoniae*. (A) Killing curve assay of WT D39 strain starting with an inoculum of 5.9×10^7^ CFU/mL in M17 media supplemented with indicated concentrations of copper sulfate and/or Compound 3. (B) Killing curve assay of WT D39 strain starting with an inoculum of 1.7×10^7^ CFU/mL in M17 media supplemented with indicated concentrations of copper sulfate and/or Compound 4. (C) Killing curve assay of WT 6303 strain starting with an inoculum of 2.3×10^7^ CFU/mL in M17 media supplemented with indicated concentrations of copper sulfate and/or Compound 3. (D) Killing curve assay of WT 6303 strain starting with an inoculum of 2.7×10^7^ CFU/mL in M17 media supplemented with indicated concentrations of copper sulfate and/or Compound 4. All bars indicate mean ± SD with n = 8-9 across 3 independent replicates. Statistical differences were measured by one way ANOVA; ns = not significant, *p < 0.05, **p < 0.01, ***p < 0.001, and ****p < 0.0001.

**Figure 8.**
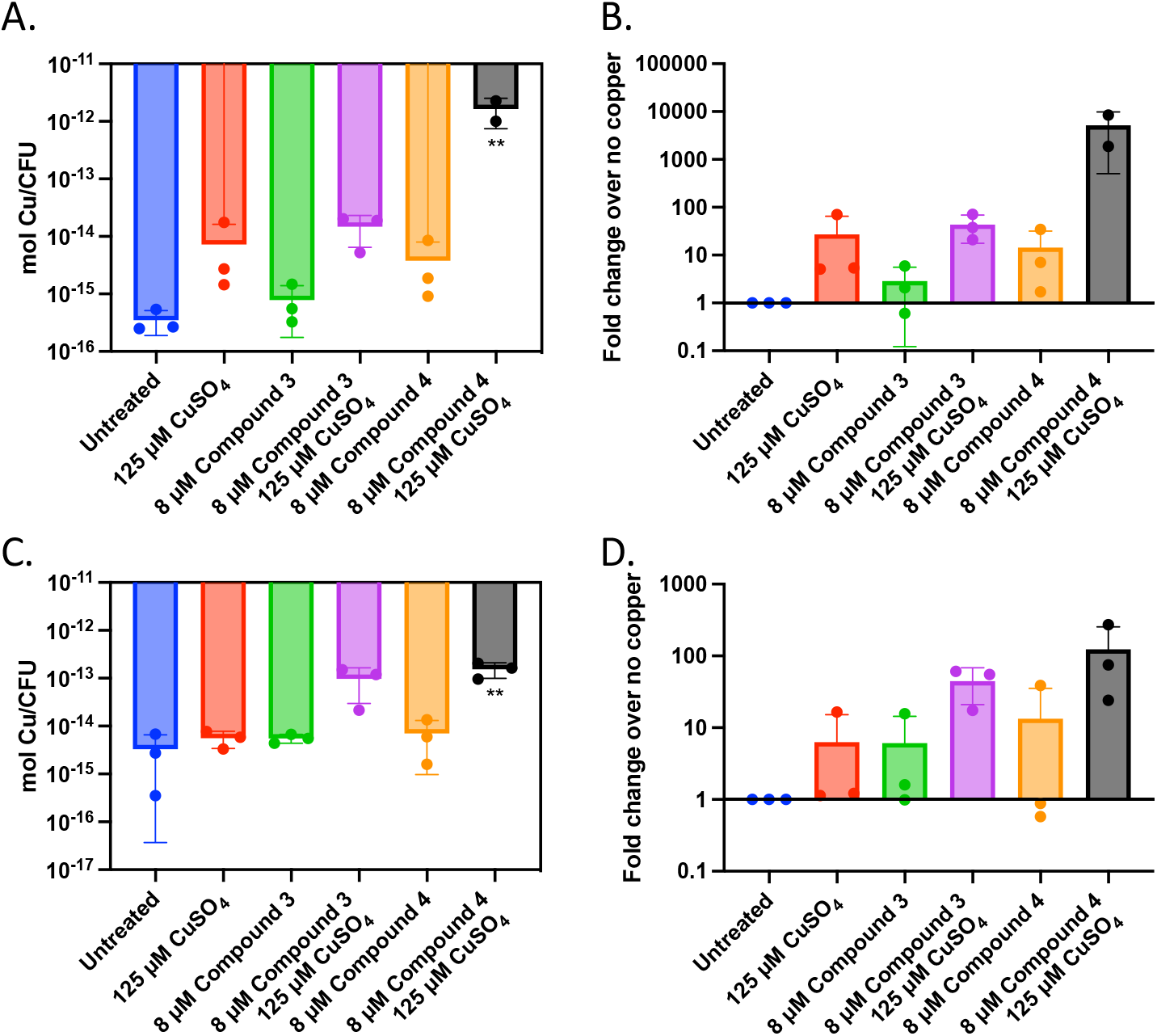
Intracellular copper concentration of D39 and Type 3 after treatment with compounds and copper. Exponentially growing D39 (A and B) or Type 3 (C and D) bacteria were treated with the indicated compound and/or copper or left untreated for 30 minutes. ICP-OES was used to measure intracellular copper. (A) and (C) represent the moles of copper per CFU while (B) and (D) represent the fold change over the respective strains with nothing added. Experiments were performed in triplicate with statistical significance determined by an Ordinary one-way ANOVA; **p < 0.01. Each point represents 6 individual sample replicates.

## Discussion

In this study, we sought to test the antimicrobial efficacy of compounds derived from *N,N*-dimethyldithiocarbamate (DMDC). We previously identified DMDC as a potent copper-dependent antibiotic against *Streptococcus pneumoniae, Staphylococcus aureus, Coccidioides immitis*, and *Schistosoma mansoni* (18). Compound 3 and Compound 4 showed *in vitro* inhibition in growth curves against TIGR4 while Compound 1, Compound 2, and Compound 5 had no effect (Figure 1 and Table 1). Further, Compounds 3 and 4 had seemingly increased killing capacity after 30 minutes compared to DMDC, which didn’t start showing killing until the 60 minute time point, thus indicating potential advancement in modifying dithiocarbamates as a copper dependent antimicrobial (21). After identifying that these DMDC derivatives could force copper into the bacteria and chelate copper, we observed a reduced bacterial burden during a murine respiratory infection using Compound 3 (Figure 2, 3, 4, 5, 7, 8). We showed that the combination of Compound 3 and copper made the pneumococcus more susceptible to activated macrophage mediated killing (Figure 6). We also tested and confirmed that Compound 3 and Compound 4 were bactericidal against the D39 strain (which has a Type 2 capsule) and against the WT Type 3 strain (ATCC® 6303™) (Figure 8). Compound 3 and Compound 4 displayed copper-dependent toxicity as an antibiotic against multiple strains of *S. pneumoniae*.

It has been shown in previous studies utilizing murine models of infection that the concentration of copper increases within the blood and lungs up to four-fold (29). Contributing to the increase in copper concentration within the lungs and blood is likely the upregulation of the acute-phase reactant ceruloplasmin, a copper-containing protein (30). With this increase in copper within the blood, it follows that the *in vivo* administration of dithiocarbamates such as Compound 3 and DMDC may be able to mimic the *in vitro* copper-dependent antimicrobial effect. Further investigation into the dithiocarbamates mechanism(s) of action are areas of active investigation.

Investigation of the use of compounds in the dithiocarbamate class has grown over the last few years. The anionic CS_2_ group characteristic of dithiocarbamates facilitates complex formation with transition metal cations and enzyme inhibitory activity (31). Recent advancements have shown the activity of dithiocarbamate derivatives as potent anticancer, antifungal, anti-neurodegenerative, and anti-inflammatory drugs (32). Potent anticancer dithiocarbamates have been identified with a variety of mechanisms of action, including DNA intercalation, inhibition of DNA topoisomerase I and II, and inhibition of other essential kinases (33). Several studies have found diethyldithiocarbamate (DEDC) to be a potent antifungal, antibiotic, and antiparasitic agent (17, 34, 35). One potential mechanism of action by which dithiocarbamate compounds can execute antibiotic activity is through transporting divalent metal ions into bacterial cells. In a study by Lanke *et al*., the authors show that a dithiocarbamate compound (pyrrolidine dithiocarbamate) causes an influx of Zn^2+^ ions into HeLa cells as a mechanism of action for its zinc-dependent antiviral activity against picornavirus (36). We hypothesize that DMDC, Compound 3, and Compound 4 exhibit similar functions, aiding transport of divalent copper ions into the bacterial cell. As an influx in intra-bacterial copper ions builds up due to dithiocarbamates shuttling them into the bacterium, mismetallation can cause bacterial enzymes to function at lower rates and lead to bacterial death.

Overall, we present data from a small-molecule screen of compounds derived from DMDC showing that Compound 3 and Compound 4 are potent copper-dependent antibiotics against several strains of *S. pneumoniae*. In addition, we show that both compounds exhibit zinc-dependent toxicity, and one of the compounds, Compound 3, exhibits potent *in vivo* efficacy as an antibiotic for aiding murine *S. pneumoniae* clearance. Further investigation into dithiocarbamate derivatives warrants further study for developing novel antimicrobials in the fight against antimicrobial-resistant pathogens.

## Materials and Methods

### Synthesis of DMDC analogs

The general synthetic procedure used was with a solution of KOH (56 mg, 1.00 mmol) in EtOH (10 mL) was cooled to 0 °C, then morpholine (87 µL, 1.00 mmol) and CS_2_ (151 µL, 2.50 mmol) were added to the solution successively. The resulting mixture was stirred at room temperature for 2 h, and then the solvent was reduced under vacuum. The first compound, Compound 1 (Dipotassium piperazine-1,4-dicarbodithiate, LRS01-057), was synthesized. ^1^H NMR (400 MHz, DMSO-*d*_6_) δ 4.34 – 4.22 (m, 4H), 3.51 – 3.39 (m, 4H). ^13^C NMR (100 MHz, DMSO-*d*_6_) δ 215.02, 66.58, 50.01.

Following the general procedure, the title compound was synthesized in 57% as a white solid to Compound 2 (Sodium 4-(*p*-tolyl)piperazine-1-carbodithioate, LRS01-084) ^1^H NMR (400 MHz, DMSO-*d*_6_) δ 4.20 (s, 8H). ^13^C NMR (100 MHz, DMSO-*d*_6_) δ 214.14, 49.51.

Following the general procedure using NaOH as a base, the title compound was synthesized in 83% as a light yellow solid to synthesize Compound 3 (Sodium *N*-benzyl-*N*-methyldithiocarbamate, LRS01-075) ^1^H NMR (400 MHz, DMSO-*d*_6_) δ 6.98 (d, *J* = 8.1 Hz, 2H), 6.80 (d, *J* = 8.2 Hz, 2H), 4.41 (t, *J* = 5.1 Hz, 4H), 2.96 (t, *J* = 5.1 Hz, 4H), 2.16 (s, 3H). ^13^C NMR (100 MHz, DMSO-*d*_6_) δ 214.73, 149.41, 129.78, 127.97, 116.15, 49.25, 49.09, 20.49.

Following the general procedure using NaOH as a base, the title compound was synthesized in 73% as a white solid for Compound 4 (Sodium *N*-allyl-*N*-methyldithiocarbamate, LRS01-077). ^1^H NMR (400 MHz, DMSO-*d*_6_) δ 7.28 – 7.19 (m, 4H), 7.19 – 7.12 (m, 1H), 5.43 (s, 2H), 3.24 (s, 3H). ^13^C NMR (100 MHz, DMSO-*d*_6_) δ 215.70, 139.58, 128.42, 127.66, 126.75, 57.85, 40.99.

Following the general procedure using NaOH as a base, the title compound was synthesized in 56% as a white solid to Compound 5 (Sodium ((*2S,3S*)-1-ethoxy-3-methyl-1-oxopentan-2-yl)carbamodithioate, LRS01-072). ^1^H NMR (400 MHz, DMSO-*d*_6_) δ 5.83 – 5.70 (m, 1H), 5.08 – 5.02 (m, 1H), 5.02 – 4.97 (m, 1H), 4.73 (d, *J* = 5.8 Hz, 2H), 3.25 (s, 3H). ^13^C NMR (101 MHz, DMSO-*d*_6_) δ 214.76, 135.06, 116.16, 57.57, 40.88.

### Bacterial Culture

M17 media (M17) (BD Difco, USA) was prepared according to the manufacturer’s instructions. Briefly, 37.25 g of powder was suspended in 950 mL of Milli-Q grade water (≥18.0 MΩ cm^-1^) and autoclaved at 121 ºC for 15 minutes before cooling to 50 ºC and adding 50 mL of a sterile 10% lactose solution. Tryptic Soy Agar (TSA) (Hardy Diagnostics, USA) was dissolved in Milli-Q water and autoclaved. After cooling autoclaved TSA, 5% defibrillated sheep’s blood (HemoStat Laboratories) of final volume and 20 μg/mL neomycin was added to the solution. These plates (blood agar plates – BAP) were used for routine culture on solid media and “killing curve” serial dilution CFU counting. Bacteria from freshly streaked plates were placed into M17 and grown at 37 °C in 5% CO_2_, to an optical density (OD or OD_600_) of 0.125 for growth curve assays and to an OD of ∼0.300 for killing curve assays. To prepare working stocks of viable *S. pneumoniae*, growing cultures are resuspended in fresh media +20% v/v glycerol and stored at -80 °C. Aliquot viability and CFU counts were determined before use in experiments. Glycerol stock aliquots were diluted 1:5 into M17 with indicated copper and compound concentrations for assays.

### Growth Curves

Growth curve assay as described in Menghani and Rivera *et al*. (18). Briefly, copper stock solutions at 100 mM were prepared from CuSO_4_ pentahydrate (VWR Life Sciences, USA) in Milli-Q water. Stock solutions of 100 mM Zn^2+^ were prepared from ZnSO_4_ heptahydrate (VWR Life Sciences, USA). Stock solutions of 1 mM Compound compounds were prepared in Milli-Q water from solute derived and purified by the University of Arizona Center for Drug Discovery (ACDD). Sterile, individually wrapped clear 96 well polystyrene plates (Greiner Bio-One, USA) were arranged to test a range of concentrations combinations of Cu^2+^, Zn^2+^, and compounds. Frozen aliquots of *S. pneumoniae* were thawed and diluted five-fold into fresh M17 before adding 20 µL per well into a total well volume of 200 µL (1:50 total dilution). Assay plates were loaded into a Biotek Cytation5 (Biotek, Vermont, USA) pre-equilibrated to 37 °C and 4% CO_2_. Gas control settings were modified for an elevation of 720 m according to the manufacturer’s directions. The protocol-maintained temperature and CO_2_, while measuring OD absorbance at 600 nm every 30 minutes for 16-20 hours.

### Killing Curves

Killing curve assay as described in Menghani and Rivera *et al* (18). Aliquots of *S. pneumoniae* were thawed and diluted ten-fold into assay conditions prepared in M17 media. Assay conditions included various concentrations of CuSO_4_, Compound compounds, Zn^2+^, and Mn^2+^. After exposure to the indicated conditions, bacteria were incubated at 37 °C in 5% CO_2_ for the stated time. Samples were serially diluted, plated on BAP, incubated overnight at 37 °C in 5% CO_2_, and counted to determine viable CFU. Colonies on each plate were counted and multiplied by the appropriate dilution factor based on which dilution it was to determine CFU/mL. For plates in which no colonies were visualized, this was deemed to be below the limit of detection (LoD) and is noted with a data point below the LoD line.

### Protein purification

As performed previously (37) BL21 cells were transformed with DsRed2-pBAD and grown overnight in a shaker incubator at 250 rpm at 37 °C on LB with ampicillin 150 µg/ml. In the morning, the culture was centrifuged and resuspended on fresh media. Terrific broth (100 µg/ml amp) was inoculated to an optical density of 0.02. The culture was incubated at 37 °C in a shaker incubator, and growth was monitored until 0.5-0.6 optical density, at which point the culture was induced with 0.1% arabinose, and the temperature was reduced to 16 °C overnight. After 16-18 hours, the culture was centrifuged, and the pellet was collected. The pellet was resuspended in buffer (50mM TRIS pH 7.4, 500mM NaCl, 25mM imidazole, 5% glycerol) and lysed by sonication after adding protease cocktail (1% protease, PMSF, and 0.2mg/ml DNase). The lysate was centrifuged to remove debris, and the cell lysate was purified on an AKTA Pure via an affinity nickel column and buffer exchanged via a desalting column. The fractions were concentrated using a 10,000 mw CO (Millipore). The protein was left to mature overnight at room temperature. Concentration was calculated by sequenced and absorbance at 280nm. Purity was assessed by 12% SDS PAGE. The protein was stored at 4 °C and used within two weeks.

### Competition assay

We performed a relative affinity test using the reversible fluorescence quenching of DsRed2 by copper (38, 39). The following was added to a 96 well plate (Greiner bio-one 655096) in a total volume of 200 µl: 400 µM of freshly dissolved compound was added in 10 mM MOPS Ph 7.2 (chelated overnight with a Chelex 100 Biorad and filtered sterilized through 0.2 µm membrane), 20 µM DsRed2, 100 µM copper (CuSO_4_). In a Biotek Cytation5, the absorbance at 585 nm was monitored for 3 hours to establish chelation of copper by the compounds and restore DsRed2 fluorescence.

### Inductively Coupled Plasma Optical Emission Spectroscopy

Experiments were performed in triplicate. *S. pneumoniae* strains were initially cultured on M17 + 5 mM lactose and frozen at -80 °C in 20% glycerol. These glycerol stocks were used as the seed stock to inoculate 150 mL of M17 + 5 mM lactose. The bacterial culture was incubated at 37 °C under 5% CO_2_ until an OD of ∼0.400 was reached. The culture was split into the indicated treatments and control. Incubation with treatments was performed at 37 °C and 5% CO_2_ for 30 minutes. Samples were quenched in a -3°C water bath to slow metabolism, followed by two washes of cold buffer (Tris 50 mM, NaCl 150 mM, EDTA 50 mM at pH 7.6), and centrifugation 3500 x g for 10 minutes at 4 °C. Cold decanted samples were resuspended in 70% HNO_3,_ followed by overnight incubation at 65 °C. After incubation, the samples were diluted to 2.5% HNO_3_ using Milli-Q water at 18.1 MΩ. Bacterial plate counts were performed in TSA + 5% Sheep’s Blood through serial dilutions, as described above. Samples were analyzed for metal content using an iCAP PRO XDUO ICP-OES with a wavelength 324.8 nm copper. Standards were made using the iCap Series Multi-element test solution ICAP 6000 series Validator from Thermo scientific, and metal content of the washed samples was calculated using the Qtegra software.

### Animal Experiments

All mouse studies were conducted with prior approval and under the guidelines of the IACUC at the University of Arizona (IACUC protocol number 18–410, R35 GM128653). All mice were maintained in a biosafety level 2 (BSL2) facility and monitored daily for signs of moribund. Eight-week-old female BALB/cJ mice (Jackson Laboratory) were anesthetized with 3% isoflurane and intranasally infected with an inoculum of 10^7^ CFU viable *S. pneumoniae* in 25 µL of Tris-buffered saline (TBS; 50 mM Tris, 150 mM NaCl, pH 7.4). Cohort controls were given 25 µL of TBS. At At 4-, 8-, or 24-hours post-infection, mice were treated with doses of intranasal Compound 3 or Compound 4 (approximately 1.6 mg/kg) in 25 µL of TBS. Mice were sacrificed by CO_2_ asphyxiation and immediately dissected for lung and blood collection 48 hours post-infection. Lung tissue was collected into 1.5 mL tubes containing 500 mL of phosphate-buffered saline (PBS; Gibco) after a brief initial wash in 500 µL of PBS to remove any excess blood during dissection. The tissue was then homogenized and centrifuged for 30 s at 400 x g. Blood samples (5 µL volume) were placed in a 45-µL volume PBS solution with heparin (10 UI/ml). Both lung and blood samples were then serially diluted at 1:10 and plated on blood agar plates and incubated overnight at 37°C and 5% CO_2_ for growth. The resulting bacterial colonies were counted for quantification and comparison.

### Macrophage Killing Assays

J774A.1 macrophages (ATCC, USA) were maintained in a 37 °C, 5% CO_2_ incubator with Dulbecco’s Modified Eagle’s Medium (DMEM; Sigma-Aldrich, USA) containing fetal bovine serum (FBS [10%vol/vol]; Sigma-Aldrich, USA), glutamine (2 mM; Sigma-Aldrich, USA), penicillin (50 units/ml; Sigma-Aldrich, USA), streptomycin (50 mg/ml; Sigma-Aldrich, USA), and NaHCO_3_ (0.015%). Cells were grown to 90% confluence in 12-well tissue culture plates (Greiner CELLSTAR®, Greiner Bio-One, USA) in 1 mL/well of growth medium. On the morning of the experiment, macrophages were washed twice with 1 mL PBS, and resuspended in 1 mL of “Serum-Free DMEM” growth medium without antibiotics, glutamine, NaHCO_3_ or FBS but supplemented with 5 ng/mL IFN-γ (Bio Basic, USA), 400 ng/mL LPS (EMD Millipore, MilliporeSigma, USA) for “priming of macrophages” experiment. For “*post hoc* killing efficiency” experiment, macrophages were resuspended in 1 mL of Serum-Free DMEM containing 32 µM DMDC, 5 ng/mL IFN-γ, and 400 ng/mL LPS.

Macrophages were incubated at 37 °C, 5% CO_2_ for 12 hours. Glycerol stocks of TIGR4 *S. pneumoniae* kept at OD 0.3 are removed from -80 °C storage, diluted into four 15 mL conical tubes of 5 mL total M17 + Lactose containing no additives (“Untreated”), 32 µM Compound 3, 250 µM CuSO_4_, and 250 µM CuSO_4_ + 32 µM Compound 3 respectively for pre-treatment of bacteria experiment. Prior to incubation of bacteria in 37 °C, 5% CO_2_, an inoculum plate is made by serial diluting 100 µL from the no additives conical. After 15 minutes of incubation, bacteria are centrifuged at 4500 x g for 10 minutes and resuspended in DMEM without antibiotics, glutamine, NaHCO_3_ or FBS. Macrophages are removed from incubation, media is removed, washed with 1 mL PBS twice and then infected with 100 µL of *S. pneumonia*e solutions for both experiment types, corresponding to a multiplicity of infection (MOI) of 10 bacteria per macrophage. The 12-well tissue culture plates were centrifuged at 200 x g for 2 minutes to facilitate co-culturing.

Wells were then washed twice with PBS at the given time-points; each wash was followed by a 5-min incubation in a 37 °C, 5% CO_2_ incubator in DMEM containing gentamicin (50 µg/ml). Macrophages were lysed in 0.02% SDS in ddH_2_O and serially diluted to determine the counts of viable intracellular bacteria. Data were normalized to the level of killing observed for the untreated TIGR4 bacteria for each assay.

### Statistical Analysis

Statistical significance was analyzed using Student t-test (two-tailed, unpaired), two-way ANOVA, or one-way ANOVA and Dunnett’s multiple comparisons test (Prism 9.20; GraphPad Software, USA). The p values were as follows: *p < 0.05, **p < 0.01, ***p < 0.001, and ****p < 0.0001.

## Acknowledgment

This study was supported by NIH grant 1R35128653. Additionally, Sanjay V. Menghani’s training is supported by an F30 Ruth L. Kirschstein Individual Predoctoral NRSA Fellowship from NIGMS (5F30GM139246-02). DsRed2-pBAD was a gift from Michael Davidson (Addgene plasmid # 54608 ; http://n2t.net/addgene:54608 ; RRID:Addgene_54608).

